# MHCBind: A Pan- and Allele-Specific Model for Predicting Class I MHC-Peptide Binding Affinity

**DOI:** 10.64898/2026.03.20.713120

**Authors:** Navyatha Peddi, Dharani Reddy Bijjula, Sanjana Gogte, Vani Kondaparthi

**Affiliations:** Department of Computer Science and Engineering, Keshav Memorial Institute of Technology, Hyderabad, Telangana, India - 500029; Drugparadigm Research Lab, Uppal, Hyderabad, Telangana, India - 500039; Keshav Memorial Engineering College, Uppal, Hyderabad, Telangana, India - 500088

**Keywords:** Major Histocompatibility Complex, T-cells, Immunotherapies, Cross-attention, Antigens, Graph Attention Network, Convolutional Neural Network, Contact Maps, Deep Learning

## Abstract

Major Histocompatibility Complex (MHC) molecules are essential to the immune system because they bind and present peptide antigens to T cells, enabling immune recognition and response. The specificity of MHC-peptide interactions is crucial for understanding immune-related diseases, developing personalized immunotherapies, and designing effective vaccines. Current computational methods, while powerful, often rely on a single type of molecular information, usually sequence, and implicitly model the interaction between the two molecules. To address these limitations, we introduce MHC-Bind, a novel deep learning framework that captures a more comprehensive and biologically relevant view of the binding event. MHCBind’s architecture employs a dual-view feature extraction strategy for both the MHC and the peptide. A Graph Attention Network (GAT) learns topological features from predicted residue contact maps, while a parallel 1D Convolutional Neural Network (CNN) captures multi-scale patterns from sequence embeddings. These four distinct feature sets are then integrated in a cross-fusion module that uses an attention mechanism to model interactions between the two molecules. Finally, a multi-layer perceptron (MLP) regression head maps the fused interaction signature to a precise binding affinity score. In rigorous comparative benchmarks against recent variants, such as NetMHCpan, MHCFlurry, and MHCnuggets, MHCBind demonstrates superior performance, achieving a significantly lower average prediction error (RMSE: 0.1485) and a higher correlation (PCC: 0.7231) in allele-specific contexts. For pan-allele tasks, it excels at correctly ranking peptides with a superior Spearman’s Correlation (SCC: 0.7102), a crucial advantage for practical applications. The framework’s design is inherently flexible, excelling in both allele-specific and pan-allele prediction tasks.

## 1. Introduction

Major Histocompatibility Complex (MHC) molecules are essential glycoproteins [1] that present peptide antigens to T cells [2], facilitating immune recognition and response to pathogens and abnormal cells [3].

### 1.1. MHC Types and Structure

In humans, MHCs are also known as Human Leukocyte Antigen (HLA) [4]. MHC proteins are of two main types [5]: MHC Class I and MHC Class II. MHC Class I molecules are essential for immune surveillance [6]. MHC class I molecules are heterodimers composed of a non-polymorphic β2-microglobulin light chain and a highly polymorphic α-chain (heavy chain) [7]. The peptide-binding cleft is located in the α-chain and consists of two α-helices that rest on an eight-stranded β-pleated sheet. The specificity of peptide binding to MHC molecules depends on the structural and chemical properties of these grooves [8], which vary across MHC alleles because of polymorphisms [9]. This diversity makes predicting MHC-peptide binding affinity accurately very challenging. Most nucleated cells display MHC class I molecules on their surface [10]. They mainly present peptides derived from intracellular proteins, such as normal self-proteins or viral proteins produced during infection. This process is part of the antigen processing “endogenous pathway”. CD8^+^ Cytotoxic T Lymphocytes (CTLs) recognize peptides attached to MHC class I molecules [11]. When these peptides are identified, CTLs can destroy infected or cancerous cells by inducing apoptosis in the target cell [12]. Dysregulation of MHC Class I-peptide binding is associated with diseases such as autoimmunity [13], cancer, viral infections, and transplant rejection, making these interactions important therapeutic targets [14].

### 1.2. MHCs Mechanism of Action

The proteasome breaks down intracellular proteins into tiny peptides. The Transporter Associated with Antigen Processing (TAP) complex, a heterodimer composed of the proteins TAP1 and TAP2, is responsible for transferring these peptides into the endoplasmic reticulum (ER). Newly produced MHC class I α-chains attach to β2-microglobulin in the ER. Tapasin is one of the chaperone proteins that help stabilize the MHC complex and promote peptide binding [15]. TAP-delivered peptides attach to the MHC class I molecules’ peptide-binding cleft. Anchor residues in the peptide that fit into pockets inside the MHC binding groove are responsible for this particular binding. A stable peptide-MHC complex is carried to the cell surface after it has formed. The T-cell receptor (TCR) of CD8+ T cells recognizes the peptide-major histocompatibility complex (MHC) on the cell surface. An immunological reaction is triggered if the TCR recognises the peptide-MHC combination.

Although experimental binding assays are considered the gold standard, their high cost, labor-intensive process, and low-throughput capacity make them impractical for thoroughly exploring the vast diversity of peptides and the extreme polymorphism of human MHC genes [16]. This has rightfully made computational prediction an essential tool. However, a significant gap remains. Most current methods rely on a single type of information-the linear amino acid sequence-and implicitly learn the rules of interaction within the hidden layers of a neural network.

To achieve a higher level of accuracy, it is recommended that a model understand the problem like a biologist: by analysing both the sequence and the structure of molecules, and by explicitly modeling how they interact. To address this, a new deep learning architecture has been developed.

To improve MHC-peptide binding affinity prediction, a cross-attention-based deep learning framework was proposed that combines sequential and structural insights for accurate affinity estimation [17]. First, a dual-modality feature extractor was developed for peptides and MHCs, leveraging Evolutionary Scale Modeling (ESM)-1b [18] embeddings to represent sequence context and contact maps for structural interactions. These features are processed through GATs to capture structural relationships and multi-scale CNNs to identify sequential patterns, ensuring thorough representation learning [19]. To model binding dynamics, a hierarchical cross-attention module was implemented that calculates interactions between MHC and peptide features across both modalities. Finally, a prediction head integrates these cross-modal [20] embeddings into affinity scores [21].

## 2. Methodology

### 2.1. Overview

The current framework is based on the principle that more accurate predictions result from a more precise representation of the data. It has three main stages: (1) parallel, dual-view feature extraction for both the MHC and peptide; (2) a cross-fusion module to model their interaction; and (3) a regression head to estimate the final binding affinity.

MHCBind’s architecture features four parallel feature extractors, as shown in Figure 1. (A) and (B) The MHC and peptide perform a GAT process on each contact map (Graph View) and a 1D CNN on sequence embeddings (Sequence View). (C) The four resulting feature sets are then fed into the cross-fusion module, which uses cross-attention to model intermolecular interactions. (D) A final MLP [22] combines the fused representations to predict a binding affinity score.

**Figure 1.**
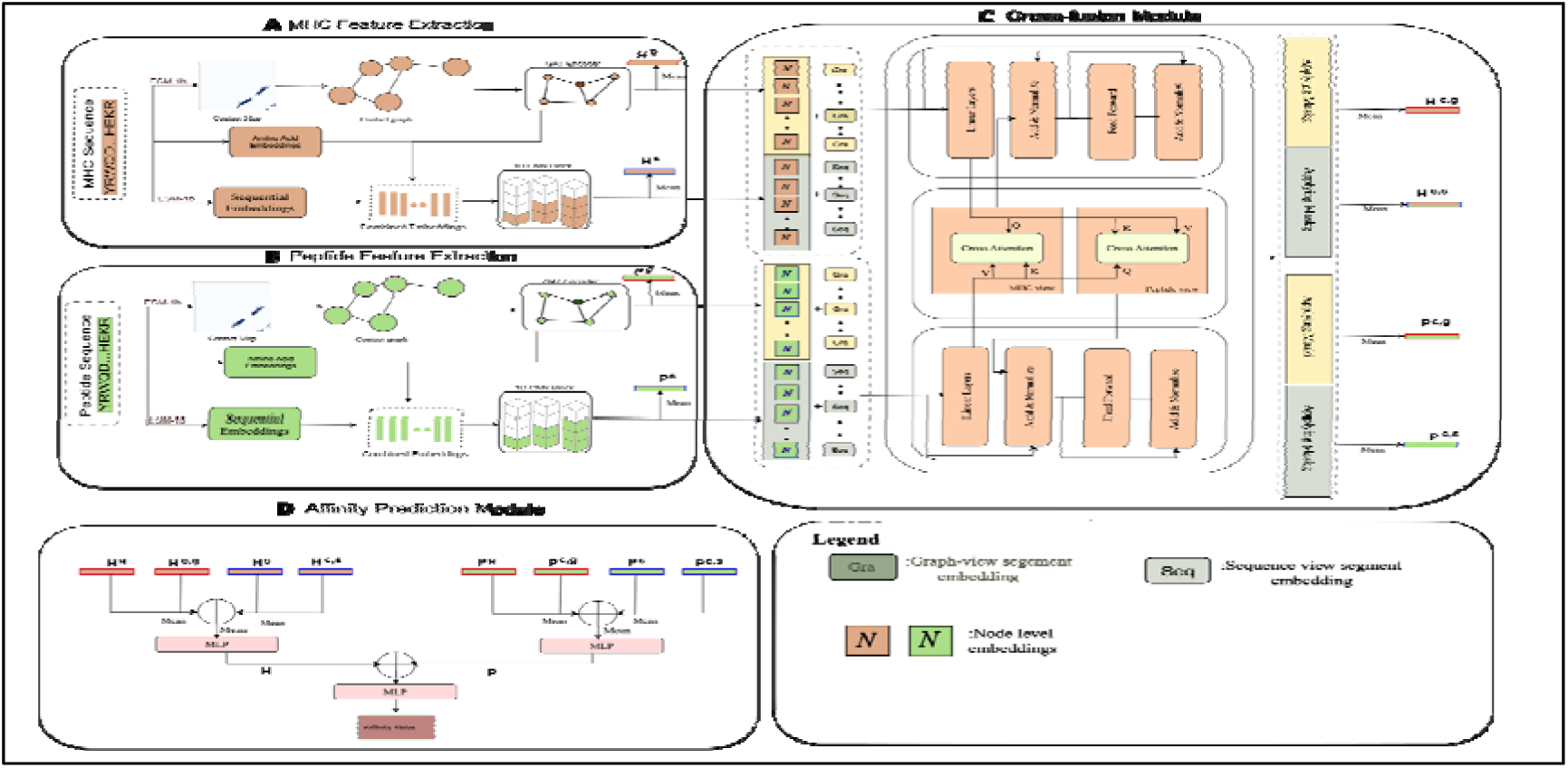
Overview of the MHCBind framework. Sections A and B depict dual-view feature extraction modules for the MHC and peptide, which generate both graph-based and sequence-based representations for each molecule separately. Section C illustrates the cross-fusion module, designed to integrate MHC and peptide information from these different views and create interaction-aware embeddings. Section D presents the final affinity prediction module, which combines various molecular representations and predicts the final MHC-peptide binding affinity score.

### 2.2. Dual-View Feature Extraction

To develop a comprehensive molecular profile, both the MHC and the peptide were processed through two separate yet parallel neural network pathways. Each pathway used initial embedding and contact map predictions [23] derived from the powerful ESM-1b protein language model [24]. ESM-1b, trained on millions of protein sequences, provides rich embedding that captures key peptides and MHC traits. It strikes a balance between power and practicality very well compared to other protein language models [25].

#### 2.2.1. Graph View using Graph Attention Network (GAT)

To analyse the 3D structural interactions of MHC molecules, the essential step is to generate contact maps [26] using the ESM-1b model, which predicts residue-residue interactions [27] based on spatial proximity [28]. For a given molecule, the amino acid sequence is first converted into a set of node features [29], □∈*R^N^*^×^*^d^*, through an embedding layer. A molecular graph is then constructed where these residues serve as nodes (v_i_), and predicted spatial contacts are represented by the adjacency matrix A. This method encodes the structural topology [30] of the MHC molecule as a graph, with residues as nodes and contacts as edges. The graph is processed through a stack of GAT layers. For a residue i with feature vector *ci* in layer l, the attention coefficient a_in_ between it and a neighbouring residue n is calculated as:

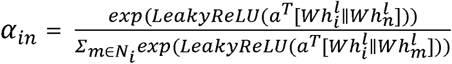

Where LeakyReLU is the activation function, W is a trainable weight matrix, a is a learnable attention vector, || denotes concatenation, and *N_i_* is the set of neighbors of node i. An aggregation step with a ReLU activation function then computes the updated feature vector *h_i_^g,l^*^+1^ for the graph view:

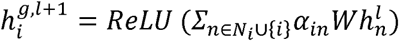

This process creates a strong representation for all residues in the molecule, labeled as *H^g^* for the MHC and *P^g^* for the peptide.

#### 2.2.2. MHC Sequence Information

This pathway emphasizes key patterns in the linear sequence. The input features, *X_seq_*, include a trainable amino acid embedding combined with pre-trained ESM 1-b embeddings. These features are fed into a 1D CNN block with multiple kernel sizes (k ∈ [3]). The convolution operation between an input sequence I and a kernel K of size k is:

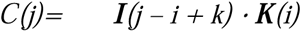

After summing the outputs of the three 1D CNN layers, a two-layer MLP projection head is employed to produce the final sequence-based feature vector, □.

Where _1_ and _2_ are trainable parameters of the linear layers, this produces the final sequence-view representations, ^s^ for the MHC and ^s^ for the peptide.

### 2.3. Cross-Fusion Interaction Module

The core of MHCBind’s framework is the Cross-Fusion Interaction Module. Instead of simply concatenating features, this module makes sure that the representations of the MHC and peptide mutually inform each other through an iterative process. It explicitly models the binding event using a Transformer-style cross-attention mechanism [31].

The process begins by preparing the inputs from the dual-view feature extraction stage (*H^s^,H^g^,P^s^,P^g^*). Before interaction can be modeled, the framework first enriches these feature sets by adding a learnable modality embedding [32, 33], allowing the model to distinguish between information derived from the sequence and the graph topology. This is achieved by adding a distinct learned vector to each feature set (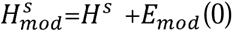 and 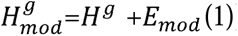), similarly for the peptide representations *P^s^,P^g^*). Following this step, the modality-aware features for each molecule are concatenated into unified input tensors,

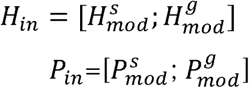

These two tensors, *H_in_* and *P_in_*, serve as the initial input to the Cross-Fusion Block.

The Cross-Fusion module facilitates bidirectional queries. The cross-attention output for the MHC view, *CrossHead^H^*, is determined as:

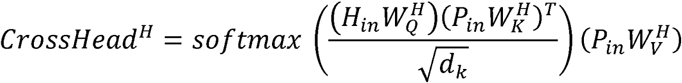

Where 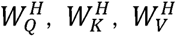 are learnable projection matrices for the Query, Key, and Value, respectively, and *d_k_* is the feature dimension. Here, the MHC acts as the “query” to attend to the peptide, which provides the “keys” and “values”. A parallel computation is performed for the peptide view, *CrossHead^p^*, where the peptide queries the MHC. This attention output is immediately integrated back with the original input via a residual connection and layer normalization. Following this, the normalized output is passed through a position-wise feed-forward Network to introduce non-linearity.

The module produces four distinct, interaction-aware output tensors: the contextualized graph and sequence views for the MHC (*H^c,g^,H^c,s^*) and the peptide (*P^c,g^,P^c,s^*).

### 2.4. Final Affinity Prediction (MLP Regression Head)

The strong, interaction-aware representations from the cross-fusion module are converted into a binding affinity score by a dedicated Multi-Layer Perceptron (MLP).

First, the different representations for each molecule are combined. The final MHC representation, H, and peptide representation, P, are derived using mean pooling and MLPs.

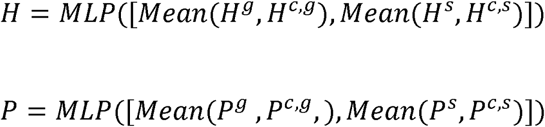

These combined vectors are concatenated and input into a final two-layer MLP to predict the binding affinity, 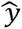:

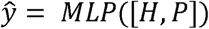

This final head simplifies the complex interaction [34] signature into a single continuous output value, mapping the learned features to the exact, quantitative scale of the experimental data.

## 3. Results and Discussion

### 3.1. Datasets and Data Curation

The foundation of any high-performance predictive model is the quality and size of its training data. A unique, high-quality dataset is curated and tailored to the specific goals of the present work, pan-allele and allele-specific models.

#### Pan-Allele dataset

For the generalizable pan-allele model, a comprehensive dataset is sourced from the Immune Epitope Database (IEDB) [35]. This top-tier resource provides an extensive and diverse collection of peptide sequences, complemented by a wide range of MHC alleles and their experimentally determined binding affinities. The breadth of this dataset was essential for training a model that could learn the fundamental rules of MHC-peptide interaction across different genetic backgrounds.

#### Allele-Specific Datasets

To develop highly specialized allele-specific models, a thorough data curation process (2022) was conducted. A unified, high-confidence dataset was created by combining information from multiple primary sources, including the IEDB and landmark high-throughput mass spectrometry (MS) [36] eluted ligand datasets [37], such as those published by Kim et al. (2014). This process involved merging data from different experimental methods, including both quantitative binding affinity assays and qualitative MS ligand identification.

### 3.2. Principled Imputation of Censored Data

A significant real-world issue in this area is handling censored measurements—values expressed as inequalities (e.g., IC50 < 50 nM or >10,000 nM). To convert this data into a format suitable for the MHCBind regression model without losing critical information, a new statistics-based imputation pipeline has been developed and implemented.

The core idea was to use the distribution of the known, quantitative measurements (=) to inform the imputation of the unknown, censored measurements (< or >). The method works in three main steps:

1. **Statistical Profiling:** Initially, a continuous numerical bin was created using all the unique inequality thresholds in the dataset. Then, for each bin, the mean and standard deviation were computed using only the known, quantitative data points (where measurement_inequality == ‘=’) that fell within that specific range.
2. **Imputation with a Truncated Normal Distribution:** For each censored data point, a new value was imputed by sampling from a truncated normal distribution. This distribution was defined by the mean and standard deviation of the known data within that point’s corresponding bin. Notably, the distribution was truncated at the bin’s boundaries, ensuring that the imputed value always respects the original inequality.
3. **Robustness Check:** To ensure statistical validity, a confidence check was performed. If a bin contained insufficient quantitative data (fewer than 10 data points), making it unreliable for statistical profiling, the imputation defaulted to a more conservative approach-uniform random sampling within the bin’s bounds.

### 3.3. Five-Fold Cross Validation

A comprehensive dataset of MHC Class I peptides was utilized, sourced from the IEDB. The final curated dataset used for our experiments consists of 1,85,985 unique peptide-MHC interactions.

To ensure a robust and unbiased assessment of our model’s performance, a 5-fold cross-validation protocol was employed across the entire dataset. This method involves partitioning the dataset into five unique segments, where each segment is sequentially held out as a validation set while the remaining four are used for training. This cycle repeats five times, ensuring each segment has a chance to serve as the validation set. A stratified K-fold for regression was specifically implemented. The results for each fold are summarized in Table 1.

**Table 1.** Metrics across all five-folds.

| Fold | MAE (↓) | RMSE (↓) | PCC (↑) | SCC (↑) |
| --- | --- | --- | --- | --- |
| 1 | **0.1288** | **0.1915** | **0.7852** | **0.7102** |
| 2 | 0.1446 | 0.2181 | 0.7409 | 0.6785 |
| 3 | 0.1619 | 0.2477 | 0.6948 | 0.6591 |
| 4 | 0.1833 | 0.2815 | 0.6095 | 0.5884 |
| 5 | 0.1624 | 0.2471 | 0.7132 | 0.6726 |

The MHCBind is benchmarked against MHCflurry 2.0 [38], focusing on four key regression metrics [39]: Pearson Correlation Coefficient (PCC), Spearman’s Correlation Coefficient (SCC), Root Mean Squared Error (RMSE), and Mean Absolute Error (MAE), which is summarized in Table 2.

**Table 2.** Overall Performance Comparison (Pan-Allele Model).

| Model | MAE (↓) | RMSE (↓) | PCC (↑) | SCC (↑) |
| --- | --- | --- | --- | --- |
| MHCBind | 0.1288 | 0.1915 | 0.7852 | **0.7102** |
| MHCflurry 2.0 | **0.1225** | **0.1714** | **0.7865** | 0.7026 |

The overall performance comparison of the Pan-Allele Model is summarized in Table 2, with numbers in bold representing the respective values.

The 5-fold cross-validation results confirm the robust performance of MHCBind’s pan-allele model, as summarized in Table 1.

It achieves a strong Pearson Correlation, peaking at 0.7852, and demonstrates excellent ranking capability with a top Spearman’s correlation of 0.7102. The consistent scores across the folds emphasize the model’s reliability and stable predictive power.

As the results indicate, MHCflurry 2.0 demonstrates a marginal advantage in metrics related to precise value prediction, achieving slightly lower RMSE and MAE. However, MHCBind achieves a significantly higher SCC. This performance trade-off is visually represented in Figure 2.

**Figure 2.**
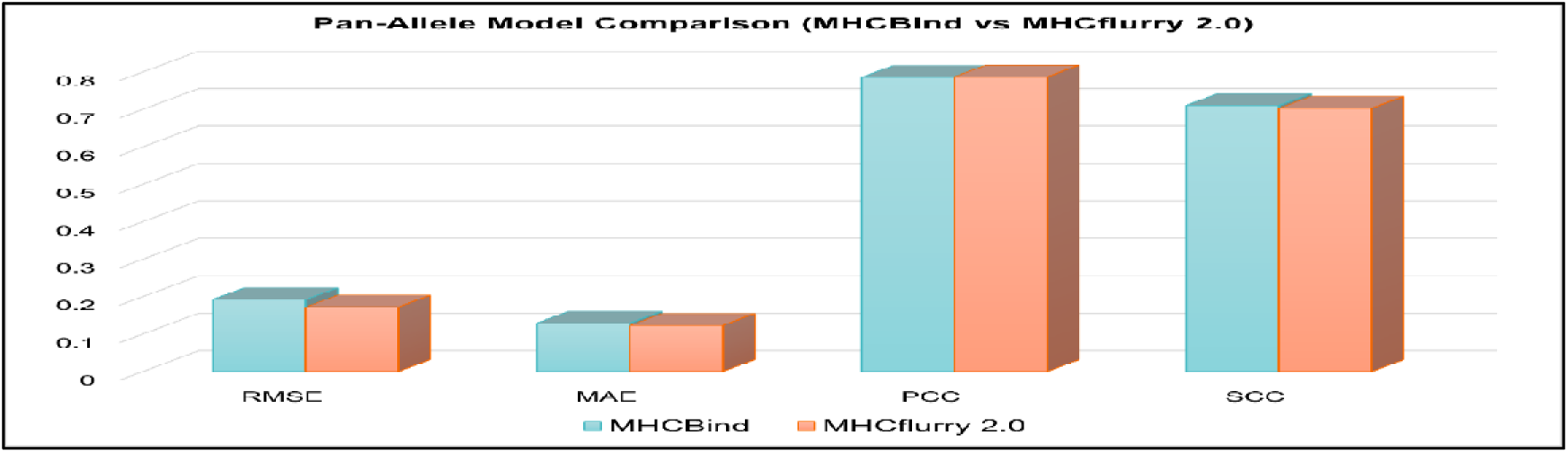
A Pan-allele Model comparison. A comparison of MHCBind against the recent variant model, MHCflurry 2.0, on four key performance metrics.

### 3.4. Allele-Specific Performance Analysis

While overall metrics are helpful, a strong model must perform well across many individual MHC alleles. The focus is on the 26 alleles with the most data, selecting only those with more than 4,000 curated binding examples to ensure a reliable training set. The results, summarized in Tables 3 and 4, show that MHC-Bind consistently outperforms current conventional methods, including NetMHCpan 4.0 [40], MHCflurry 2.0 [38], and MHCnuggets [41], as illustrated in Figures 3 and 4. To highlight this, a detailed allele-specific analysis is conducted. The performance of MHC-Bind compared to state-of-the-art models for each of the four key metrics across various MHC alleles is displayed as line plots in Figure 5, covering RMSE [42], MAE [43], PCC [44], and SCC [45]. MHC-Bind consistently surpasses existing methods across a broad range of alleles.

**Figure 3.**
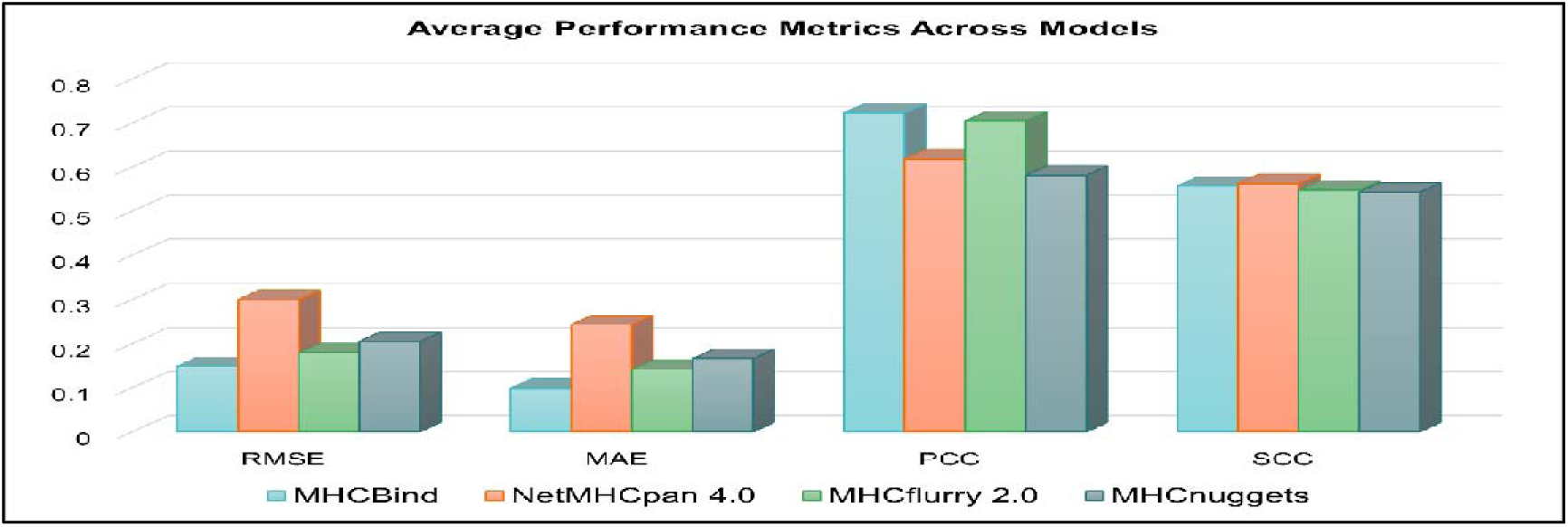
Overall performance (Average Metrics). A comparison of MHCBind against the state-of-the-art models, showing the average RMSE, MAE, PCC, and SCC across all allele-specific models.

**Figure 4.**
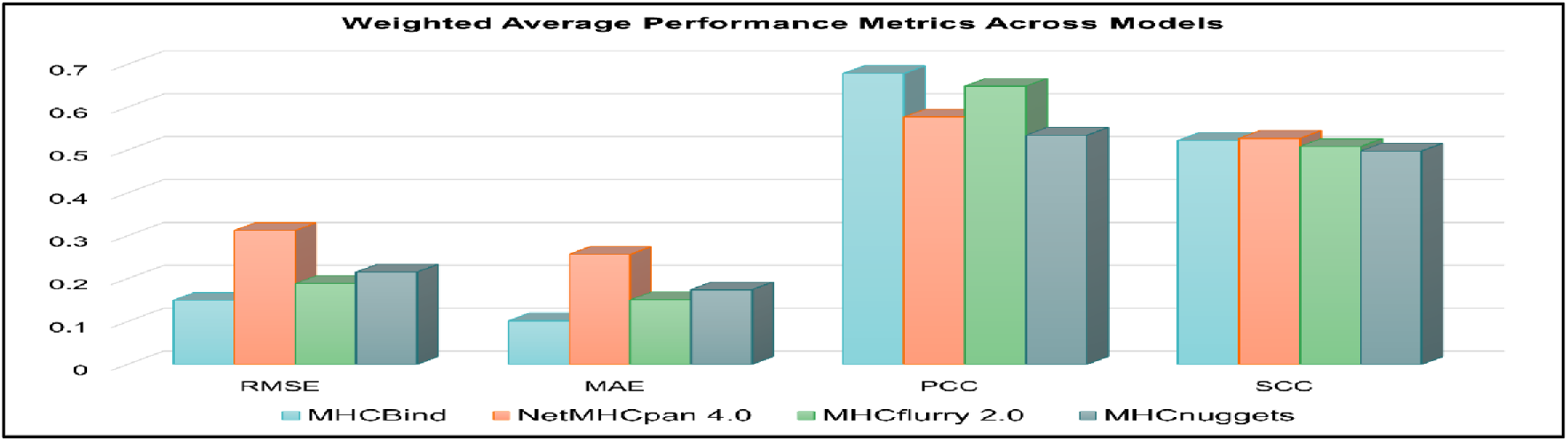
Overall Performance (Weighted Average Metrics). A performance comparison showing the weighted average of RMSE, MAE, PCC, and SCC. Metrics are weighted by the test sample size of each allele to provide a more representative score that accounts for data imbalance.

**Figure 5.**
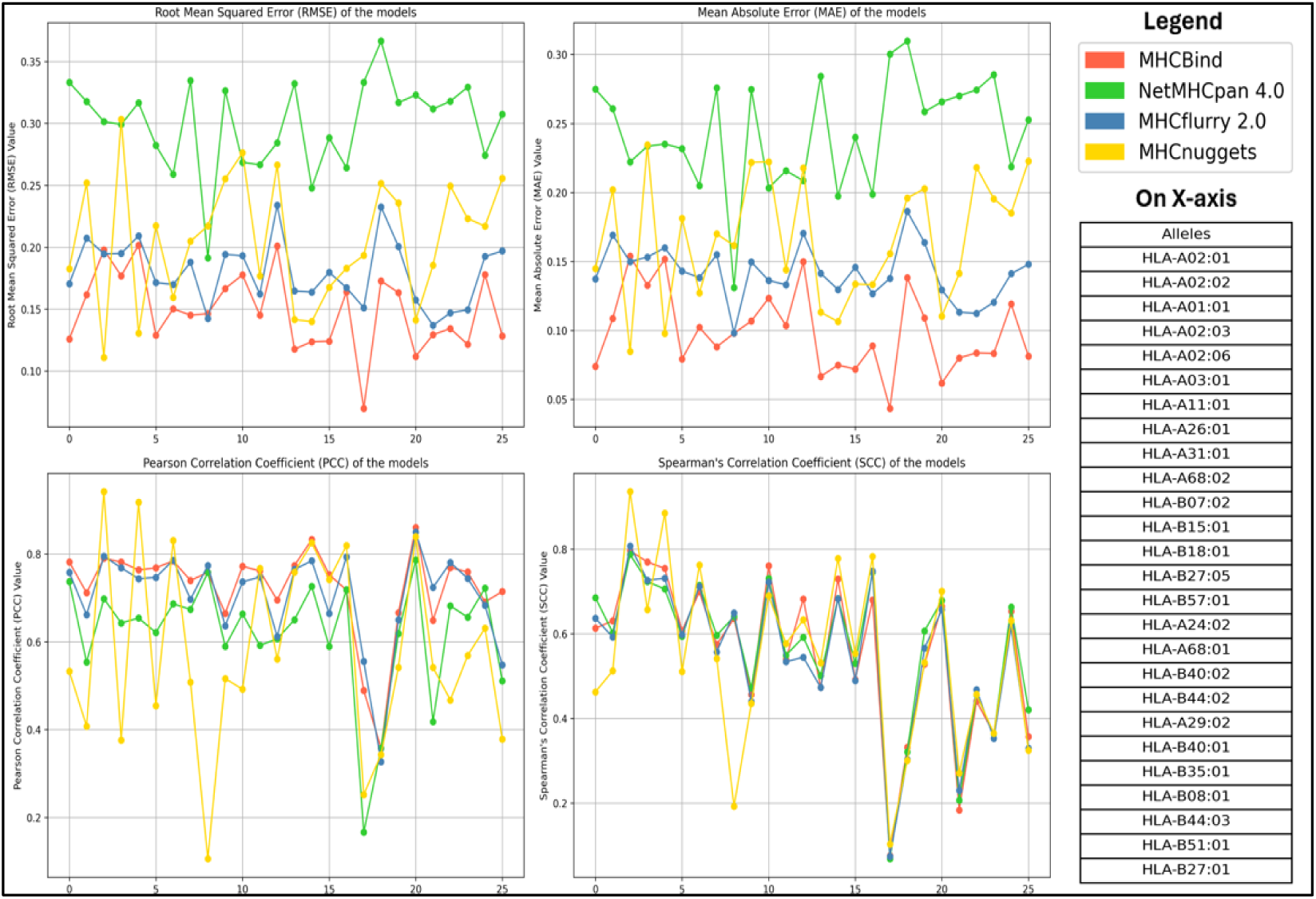
The MHCBind performance comparison. Comparison to the recent variant models across various individual MHC alleles is depicted in Figure 5. The four line plots consistently show that the MHCBind model has an advantage in RMSE, MAE, PCC, and SCC, demonstrating its strong performance across different alleles.

**Table 3.** Overall Performance Comparison (Average Metrics).

| Model | RMSE (↓) | MAE (↓) | PCC (↑) | SCC (↑) |
| --- | --- | --- | --- | --- |
| NetMHCpan4.0 | 0.2997 | 0.2433 | 0.6183 | **0.5629** |
| MHCflurry 2.0 | 0.1796 | 0.1418 | 0.7048 | 0.5476 |
| MHCnuggets | 0.2052 | 0.1662 | 0.5815 | 0.5431 |
| MHCBind | **0.1485** | **0.0990** | **0.7231** | 0.5573 |

The simple average metrics for MHCBind compared to NetMHCpan, MHCflurry, and MHCnuggets across 26 alleles are presented in Table 3.

**Table 4:** Overall Performance Comparison (Weighted Average Metrics).

| Model | RMSE (↓) | MAE (↓) | PCC (↑) | SCC (↑) |
| --- | --- | --- | --- | --- |
| NetMHCpan4.0 | 0.3120 | 0.2566 | 0.5774 | **0.5273** |
| MHCflurry 2.0 | 0.1885 | 0.1504 | 0.6493 | 0.5083 |
| MHCnuggets | 0.2153 | 0.1740 | 0.5343 | 0.4979 |
| MHCBind | **0.0229** | **0.1013** | **0.6786** | 0.5227 |

Weighted averages are calculated by multiplying each metric by its corresponding dataset size, summing these products across allele models, and then dividing by the total dataset size, thereby emphasizing the reliability of larger datasets. In contrast, simple average treats each allele-specific model’s metric equally, ignoring the dataset size. This approach to calculating weighted averages is significant and effective, as it better reflects the model’s performance on diverse dataset sizes, ensuring that metrics are fairly evaluated across allele-specific models.

The results from both the average and weighted average analyses consistently highlight the superior predictive accuracy of the MHCBind model. MHCBind achieves a notable advantage in quantitative precision, demonstrating lower RMSE and MAE, as well as PCC. This indicates our model’s enhanced capability to predict the exact binding affinity value and maintain a strong linear relationship with the experimental data.

While the SCC, which measures the rank order, is competitively close to the state-of-the-art, the MHCBind model’s primary strength lies in its quantitative accuracy. This suggests that for applications where the precise estimation of binding affinity values is the most critical factor, MHCBind offers a distinct and reliable advantage.

MHCBind’s strong performance is not due to an averaging effect, but rather because it remains consistent across individual alleles. The plots clearly demonstrate that MHCBind often achieves the highest correlation and the lowest error for various MHC types. This consistent performance across a range of alleles with diverse binding groove shapes and properties suggests that the MHCBind model is not simply memorizing allele-specific sequence motifs. Instead, it shows that the dual-view architecture has effectively learned more generalizable, underlying biophysical principles [46] of peptide binding.

## 4. Conclusion

In this study, MHCBind is introduced as a novel deep learning platform specifically designed to address the long-standing challenge of predicting MHC–peptide binding affinity. The main innovation of MHCBind is its dual-view architecture, which processes sequence-level and structural topology features simultaneously through two computational streams: a Graph Attention Network (GAT) branch that captures structural dependencies and a 1D Convolutional Neural Network (CNN) branch that extracts contextual patterns and local sequence motifs. These parallel representations are then merged via a specially designed cross-fusion module, enabling the model to construct a biophysically informed latent space that integrates both structural and sequential factors influencing binding affinity.

Our findings demonstrate that this combined strategy provides a significantly richer representation of MHC–peptide interactions than what current sequence-only models can achieve. In allele-specific prediction tasks, MHCBind sets a new standard with an average RMSE of 0.1485 and a Pearson correlation coefficient (PCC) of 0.7231, surpassing many improved baseline approaches. Importantly, in the more realistic pan-allele prediction scenario, where generalization across unseen alleles is crucial, MHCBind attains a Spearman correlation coefficient (SCC) of 0.7102, highlighting its strong ability to rank candidate peptides even in data-scarce situations.

The importance of these findings is immense and groundbreaking for computational immunology. For the first time, better prediction accuracy directly reduces experimental work by helping researchers focus only on the most promising peptide candidates for follow-up testing. This is especially important in vaccine development, cancer neoantigen discovery, and infectious disease immunotherapy, where the rapid and accurate identification of immunogenic epitopes can significantly accelerate the transition from computational screening to experimental validation. Additionally, the dual-view approach provides a modelable solution to other biomolecular interaction challenges, demonstrating how graph-based structural learning can be combined with sequence modeling to capture both local and global factors that influence interactions.

Looking ahead, MHCBind paves the way for several exciting research directions. Future developments could include protein language models for more detailed allele embeddings, attention-based interpretability modules to clarify molecular recognition mechanisms, and multi-task learning setups to predict co-related immunogenicity endpoints simultaneously. Furthermore, combining high-throughput experimental data with structure-prediction tools (e.g., AlphaFold2) may enable the creation of comprehensive systems that go from raw sequence data to immunogenicity predictions.

In summary, MHCBind not only sets a new performance benchmark for predicting MHC–peptide binding affinity but also introduces a novel modeling approach that connects biophysical understanding with deep learning. We anticipate that this work will significantly advance computational immunology research, providing practical value for epitope discovery and inspiring new methods for structure-aware biological machine learning.

## Declarations

### Ethics approval and consent to participate

Not applicable

### Consent for publication

Not applicable

## Code availability

The code is available at GitHub Link: https://github.com/drugparadigm/MHCBind/tree/main

## Author Contributions

**N.P.:** Investigation; validation; methodology; visualization; writing-original draft; formal analysis. **D.B.:** Data curation; formal analysis; methodology; validation; visualization. **S.G.:** Conceptualization; methodology; validation; visualization; software; supervision; writing-review and editing. **V.K.:** Conceptualization; methodology; writing-review and editing; supervision; formal analysis.

## Acknowledgements

We, the authors NP, DB, SG, and VK, express our sincere gratitude to the Drugparadigm Research Lab for providing the necessary facilities and infrastructure that enabled the successful completion of this work.

